# Drug dependence in cancer is exploitable by optimally constructed treatment holidays

**DOI:** 10.1101/2022.07.01.498458

**Authors:** Jeff Maltas, Katherine R. Singleton, Kris C. Wood, Kevin B. Wood

## Abstract

Recent work in cell culture models, animal models, and human patients indicates that cancers with acquired resistance to a drug can become simultaneously dependent upon the presence of that drug for survival. This drug *dependence* offers a potential avenue for improving treatments aimed at slowing resistance, yet relatively little is known about the frequency with which drug dependence arises, the mechanisms underlying that dependence, and how drug schedules might be tuned to optimally exploit drug dependence. In this work, we address these open questions using a combination of laboratory evolution, *in vitro* experiments, and simple mathematical models. First, we used laboratory evolution to select more than 100 resistant *BRAF* mutant melanoma cell lines with acquired resistance to BRAF, MEK, or ERK inhibitors. We found that nearly half of these lines exhibit drug dependence, and the dependency response is associated with EGFR-driven senescence induction, but not apoptosis, following drug withdrawal. Then, using melanoma populations with evolved resistance to the BRAF inhibitor PLX4720, we showed that drug dependence can be leveraged to dramatically reduce population growth when treatment strategies include optimally chosen drug-free “holidays”. On short timescales, the duration of these holidays depends sensitively on the composition of the population, but for sufficiently long treatments it depends only on a single dimensionless parameter (*γ*) that describes how the growth rates of each cell type depend on the different treatment environments. Experiments confirm that the optimal holiday duration changes in time–with holidays of different durations leading to optimized treatments on different timescales. Furthermore, we find that the presence of “non-dependent” resistant cells does not change the optimal treatment schedule but leads to a net increase in population size. Finally, we show that even in the absence of detailed information about the composition and growth characteristics of cellular clones within a population, a simple adaptive therapy protocol can produce near-optimal outcomes using only measurements of total population size, at least when these measurements are sufficiently frequent. As a whole, these results may provide a stepping-stone toward the eventual development of evolution-inspired treatment strategies for drug dependent cancers.

## INTRODUCTION

The advent of targeted therapies has allowed for improved outcomes in oncology. However, even the most successful targeted drugs are not often curative, especially for patients with metastatic cancers who ultimately progress with drug resistant disease (1, 2, 3, 4). Despite tremendous progress, our most successful cancer treatment strategies often fail to explicitly account for cellular heterogeneity and disease evolution. Recently, though, as metastatic drug resistance continues to evade our best clinical practices, there has been a movement toward evolution-informed therapies (5, 6, 7, 8). These therapies aim to treat cancers (and other diseases such as those caused by pathogenic bacteria, viruses, fungi, and parasites) as heterogeneous, ecologically diverse, evolving populations. For example, recent laboratory work in cancer and bacteria aims to take advantage of drug sensitivities that evolve as a result of the original treatment, a phenomenon known as “collateral sensitivity” (9, 10, 11, 12, 13, 14, 15, 16, 17, 18). More ecologically-inspired therapies take advantage of the competition for resources between distinct clonal populations, hypothesizing that more treatable clones may win-out in a competition for shared resources at a particular drug concentration (6, 19, 20, 21). Evolutionary game theory has also been leveraged to quantify how distinct clonal populations interact and develop treatments that take advantage of the “game” the population is playing (22, 23, 24, 25, 26, 27, 28, 29, 30, 31, 32). Other evolution-based strategies model evolution on fitness landscapes (33, 34, 35) or attempt to exploit drug combinations (36, 37, 38, 39, 40) or incorporate evolutionary consequences of spatial structure (41, 42, 43, 44, 45, 46, 47, 48) or dynamic environments (49, 50, 51, 52, 53, 54, 55, 56, 57) to shape evolutionary trajectories. Still, evolution-based therapies in cancer are in their infancy, as the vast majority of the existing literature consists of purely theoretical studies or clinical studies that rely largely on biological intuition (58, 59, 60, 61, 62).

Meanwhile, drug dependent (sometimes called “drug addicted”) cancers, in which a population of drug resistant cells grows faster in the presence of a drug than without it, have been identified in cell culture, animal models and human patients (63, 64, 65, 66, 67, 68, 69, 70), most often in the context of targeted therapies. While candidate mechanisms underlying drug addiction have been proposed (71, 72), it is not yet clear how an evolution-inspired therapy might optimally treat a cancer cell population that includes drug dependent cells.

In this work, we take an important step toward answering this question by investigating how intermittent therapeutic strategies, or drug holidays, may be optimally administered to mixed populations of cancer cells comprised of drug sensitive and drug resistant-dependent populations in cell culture. First, we use laboratory evolution to show that approximately half of drug resistant cells lines selected by BRAF-MEK-ERK pathway inhibitors are also drug dependent, and this dependence is associated with EGFR-driven senescence induction following drug withdrawal. Then, using a combination of proliferation assays and simple mathematical models, we show that the size of the total cell population can be dramatically reduced by intermixing drug treatments with optimally chosen drug holidays. On short timescales, the optimal time in holiday depends sensitively on the composition of the population–the ratio of sensitive to resistant cells–but on longer scales depends on a single dimensionless parameter (*γ*) that describes how the growth rates of each cell type depend on treatment environment (with or without drug). Using melanoma populations with evolved resistance to the BRAF inhibitor PLX4720, we show experimentally that the optimal holiday duration changes in time–with different holiday durations optimizing population growth on different timescales. Furthermore, we find that the presence of “non-addicted” resistant cells does not change the optimal treatment schedule but leads to a net increase in population size. These results suggest that for a range of treatment durations, a detailed knowledge of population composition is required to design optimal holidays. However, we show that in the absence of information about composition, a simple adaptive therapy protocol can produce near-optimal outcomes using only measurements of total population size, at least when these measurements are sufficiently frequent.

## RESULTS

### A subset of BRAFi-resistant, *BRAF* mutant melanoma cell lines exhibit dependence on the selecting drug

Approximately 50 percent of melanomas harbor oncogenic BRAF mutations, and these tumors are highly responsive to inhibitors of the BRAF-MEK-ERK pathway. This responsiveness, and the eventual development of acquired resistance, can be effectively modeled in BRAF mutant melanoma cell culture models, which typically exhibit exquisite sensitivity followed by long-term acquired resistance to BRAF-MEK-ERK pathway inhibitors. To characterize drug dependent resistance in laboratory populations, we experimentally selected (and STR profiled) a collection of *BRAF* mutant melanoma cell lines by exposing populations to one of three inhibitors (the BRAF inhibitor PLX4720, the MEK inhibitor AZD6244, or the ERK inhibitor VX-11E) and generating multiple pooled or clonal resistant cell lines (Materials and Methods). To characterize the response of these cell lines to drug, we plated them at low densities both in the presence and absence of the selecting drug (PLX4720, AZD6244 or VX-11E) and assessed growth by quantifying colony area. As expected, we found that parental cell lines could give rise to resistant progeny exhibiting one of two opposing responses to drug: 1) non-dependent resistance, in which resistant cells lines were more viable in drug-free (DMSO) environments than in the presence of drug, and 2) drug dependent resistance, in which resistant cells were more viable in the presence of drug (Fig 1A, Table S1).

**FIG 1.**
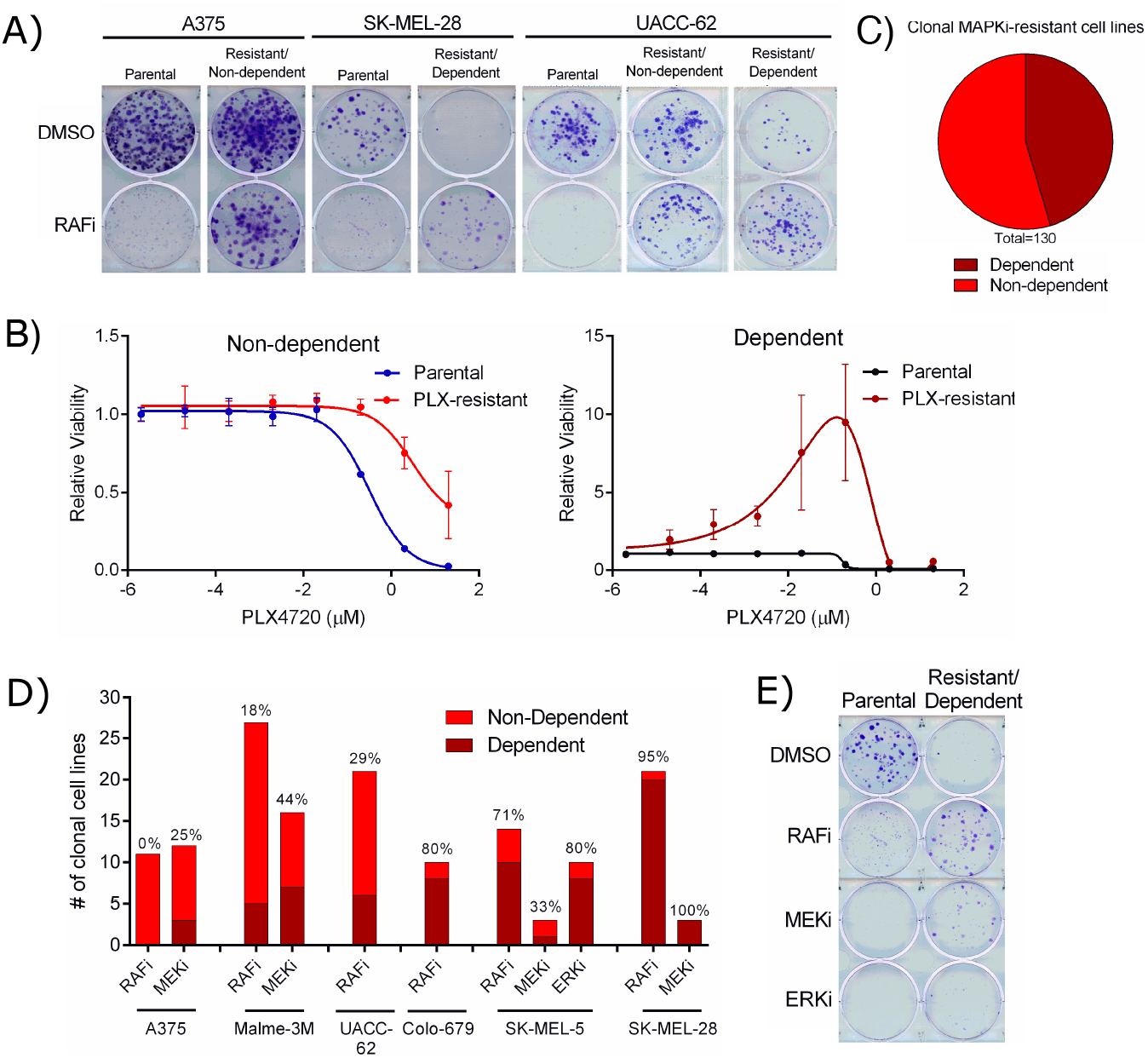
A subset of BRAF-MEK-ERKi-resistant BRAF-mutant melanoma cell lines develop dependency on the selecting inhibitor. A) The indicated cell lines were plated at a low density in normal culture conditions. Twenty-four hours later, they were treated with DMSO or 3 μM PLX4720 (RAFi) as indicated for 14 days. Resulting colonies were fixed and stained. B) A375 parental and resistant (non-dependent, left panel) or SK-MEL-28 parental and resistant (dependent, right panel) cell lines were treated with a 10-fold dilution series of PLX4720 and viability relative to DMSO treated control was determined. C) Clonal BRAF-MEK-ERKi-resistant cell lines were screened for dependency, characterized by greater colony formation in PLX4720 than in DMSO. D) As in C), but separated by cell line and selecting drug. E) SK-MEL-5 parental or SK-MEL-5 PLX4720-resistant clone 1 cell line were plated at a low density in normal culture conditions. Twenty-four hours later, they were treated with DMSO, 3 μM PLX4720 (RAFi), 3 μM AZD6244 (MEKi) or 3 μM VX-11E (ERKi) as indicated for 14 days. Resulting colonies were fixed and stained.

To quantify these phenotypic responses, we treated both parental and resistant cells with a 10-fold dilution series of PLX4720 and assayed cell viability after 3 days (Materials and Methods). While the non-dependent resistant line, a derivative of A375 (left panel), was able to survive and grow in higher doses of PLX4720 than its sensitive parental counterpart, it showed no advantage in growth in drug versus DMSO (Fig 1). Conversely, the dependent line, a derivative SK-MEL-28 (right panel), showed both a dramatic increase in growth and survival as the PLX4720 dose was increased as well as the ability to survive in higher doses than the corresponding parental line (Fig 1). Overall, we found that slightly less than half (45.4 percent) of all resistant cell lines were classified as drug dependent (percent change < −0.1) (Fig 1C and Table S1). Certain cell lines (e.g. SK-MEL-28 and SK-MEL-5) nearly always gave rise to dependent resistant cell lines, while others (A375) rarely did (Fig 1D). Additionally, inhibitors of ‘lower’ nodes of the BRAF-MEK-ERK pathway appeared to be slightly more likely to produce dependency upon resistance (Fig 1D). We also observed that cell lines with a dependency on one inhibitor in the pathway, such as PLX4720, also often displayed dependency on inhibitors of the other nodes, albeit with less resistance (Fig 1E).

### The dependency response is associated with senescence and quiescence, not Apoptosis

To investigate the mechanism of the observed drug resistance, we withdrew PLX4720 from one resistant non-dependent line (derived from A375) and two resistant dependent lines (derived from SK-MEL-5 and derived from SK-MEL-28) and measured the induction of markers of two terminal forms of growth inhibition associated with drug dependence: senescence (PAI-1) and apoptosis (cleaved PARP). We found that while we never observed significant expression of cleaved PARP, both of the resistant dependent lines strongly induced PAI-1 expression by 48 hours post drug withdrawal (Fig 2A). Using Annexin V staining as an additional measure of apoptosis, we observed less apoptosis in the resistant non-dependent cells upon drug withdrawal, concurrent with their increased growth rate off drug. However, we observed no significant change in Annexin V staining in the resistant dependent cells on drug withdrawal (Fig 2B). In agreement with these results, we observed an increase in G1 phase cells and a decrease in S and G2 phase cells on drug withdrawal in the resistant dependent cells with the opposite result in resistant non-dependent cells (Fig 2C). Additionally, resistant dependent cells showed an increase in senescence associated β-galactasidase (SA β-gal) activity on drug withdrawal while resistant non-dependent cells did not (Fig 2D). We next wished to establish whether the observed dependency related senescence was occurring by the p53 or Rb pathway (73). To investigate this, we withdrew PLX4720 from the cell types described above and immunoblotted for p53 or p16 (Rb pathway) expression. We found that while the resistant non-dependent cells displayed no detectable expression of either protein, the resistant dependent cell line, SK-MEL-5, began to express p53 at 48 hours with no expression of p16 and SK-MEL-28 began to express p16 at 48 hours while actually displaying a decrease in p53 expression (Fig 2E). This cell line has been shown to have a p53 mutation, indicating that p53 could be having a senescence suppressive function in this case(74). These results indicate that resistant dependent cells might employ different mechanisms to induce senescence on drug withdrawal.

**FIG 2.**
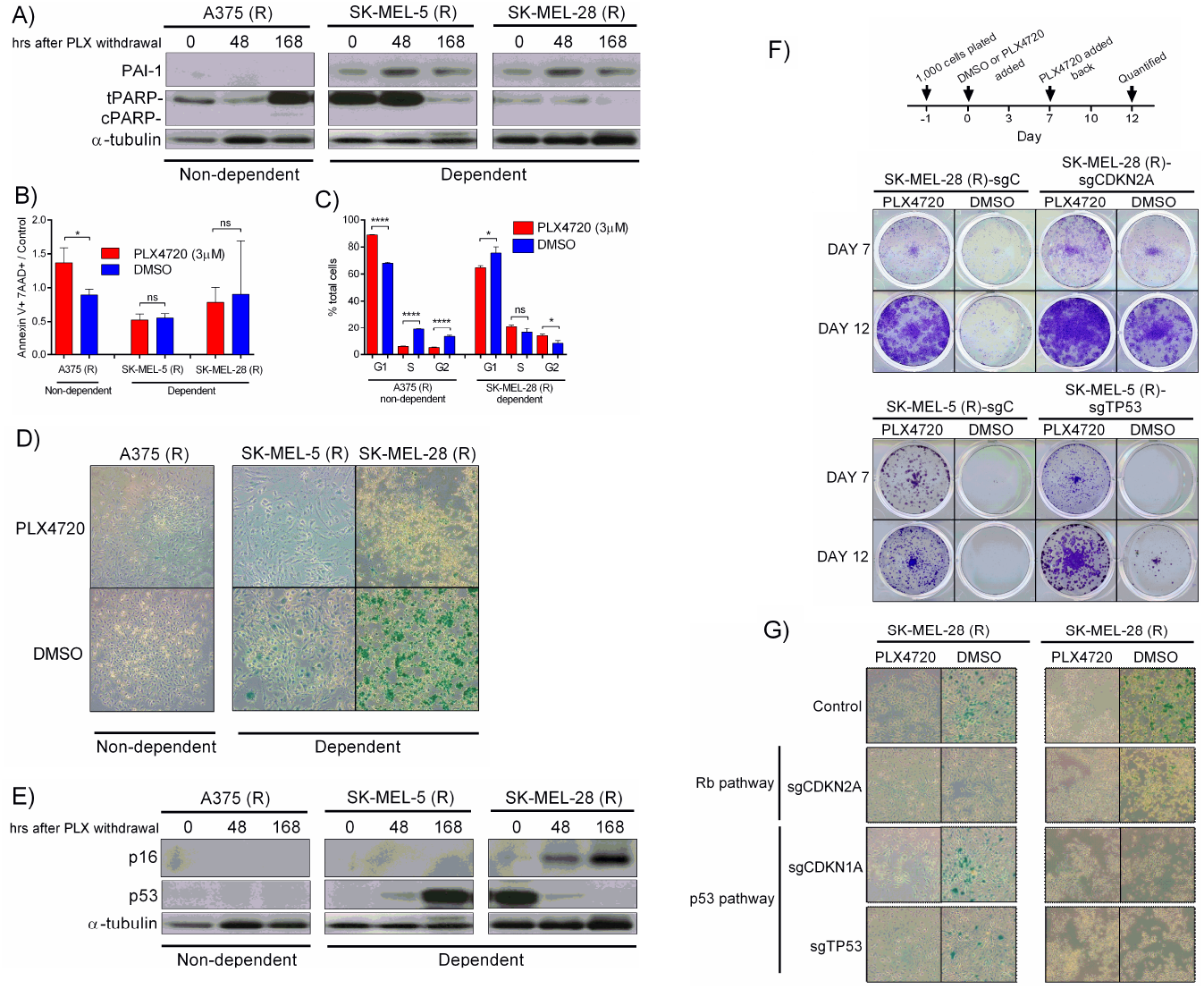
Dependent cell lines exhibit evidence of senescence upon drug withdrawal and dependency can be blocked by knockdown of senescence pathway genes. A) A375, SK-MEL-5 and SK-MEL-28 PLX4720-resistant lines were treated with PLX4720 withdrawal for the indicated time and immunoblotted for PAI-1, total and cleaved PARP and α-tubulin as a loading control. B) A375, SK-MEL-5 and SK-MEL-28 PLX4720-resistant lines were treated with PLX4720 withdrawal for 3 days and stained for annexin V and 7AAD as a measure of apoptosis. Data is normalized to cells treated with PLX4720. C) A375 and SK-MEL-28 PLX4720-resistant lines were treated with DMSO or PLX4720 for 3 days and cell cycle analysis was performed as described in the Material and Methods. The percent of total cells in each phase is shown. D) A375, SK-MEL-5 and SK-MEL-28 PLX4720-resistant lines were plated at low density and treated with PLX4720 or DMSO the following day. After 14 days, cells were stained for SA β-galactasidase activity as described in Materials and Methods. E) A375, SK-MEL-5 and SK-MEL-28 PLX4720-resistant lines were treated with PLX4720 withdrawal for the indicated time and immunoblotted for p16, p53 and α-tubulin as a loading control. F) SK-MEL-28 PLX-resistant lines expressing sgControl or sgCDKN2A and SK-MEL-5 PLX-resistant lines expressing sgControl or sgTP53 were plated at low density and treated according to the schematic (left panel). On day 7 and 12, cells were fixed and stained. G) SK-MEL-28 and SK-MEL-5 PLX-resistant lines expressing sgControl, sgCDKN2A, sgCDKN1A or sgTP53 were treated and stained for SA β-galactasidase activity as in D).

### Blocking senescence partially prevents the dependency phenotype

We next asked whether dependency could be blocked by blocking senescence or whether other mechanisms played a partial role in the dependency phenotype. To address this, we knocked out either CDKN2A (p16) in the resistant SK-MEL-28 cells that had previously been shown to employ the Rb arm of the senescence pathway or TP53 (p53) in the resistant SK-MEL-5 cells that had been shown to employ the p53 arm of the senescence pathway (Figure S1A). We were surprised to observe that even in these knockout cells, 7 days of drug withdrawal still was capable of inducing dependency (Fig 2F, top panels). However, when drug was reintroduced for 5 additional days after the drug withdrawal, the knockout, but not the control, cells were able to grow out (Fig 2F, bottom panels). The change in growth rate between the drug withdrawal and drug readdition periods is shown in Figures S1B-C. These results indicate that dependent cells preferentially utilize senescence on drug withdrawal, but that the dependent response is robust enough to employ quiescence when senescence pathways are blocked. In agreement with these hypothesis, we found no evidence of increased SA β-gal activity on drug withdrawal when either p16 (SK-MEL-28 resistant) or p53 (SK-MEL-5 resistant) was knocked out as compared to the increase in the control cells (Fig 2G), while the knockouts did not significantly change the proportion of cells in each phase of the cell cycle (Figure S1D).

### In dependent cell lines, overexpressed EGFR drives increased phospho-ERK and senescence on drug withdrawal

We next wished to establish the upstream signaling events that lead to increased p53 or p16 expression and senescence induction upon PLX4720 withdrawal in resistant dependent cell lines. Previous groups have shown a link between drug scheduling benefits and increases in phosphorylated ERK on drug withdrawal(66, 65, 71, 72). To examine whether this was occurring and to better establish the mechanisms leading to dependency, we screened a panel of resistant non-dependent and dependent cell lines for phospho-ERK when cultured in PLX4720 and when PLX4720 had been withdrawn for 72 hours. We observed a much larger in-crease in phospho-ERK among dependent lines than non-dependent lines (Fig 3A and S2A). Given the established role of EGFR in driving ERK reactivation in the setting of BRAF inhibition, we next hypothesized that increases in EGFR expression could be driving both the resistance and dependency associated increase in phospho-ERK. Indeed, we found large increases in EGFR expression between parental and resistant dependent lines, but not between parental and resistant non-dependent lines (Fig 3B and S2B). There was a strong relationship between EGFR induction and drug dependence in resistant cell lines (Fig 3C). Additionally, we could block loss of cell growth, increases in phospho-ERK and PAI-1 on PLX4720 withdrawal with the addition of the EGFR inhibitor, gefitinib (Fig 3D-E and S2C). Conversely, the combination of PLX4720 and gefitinib re-sensitized the resistant cells to PLX4720, indicating that increased EGFR expression leads to both resistance and dependency (Fig 3E).

**FIG 3.**
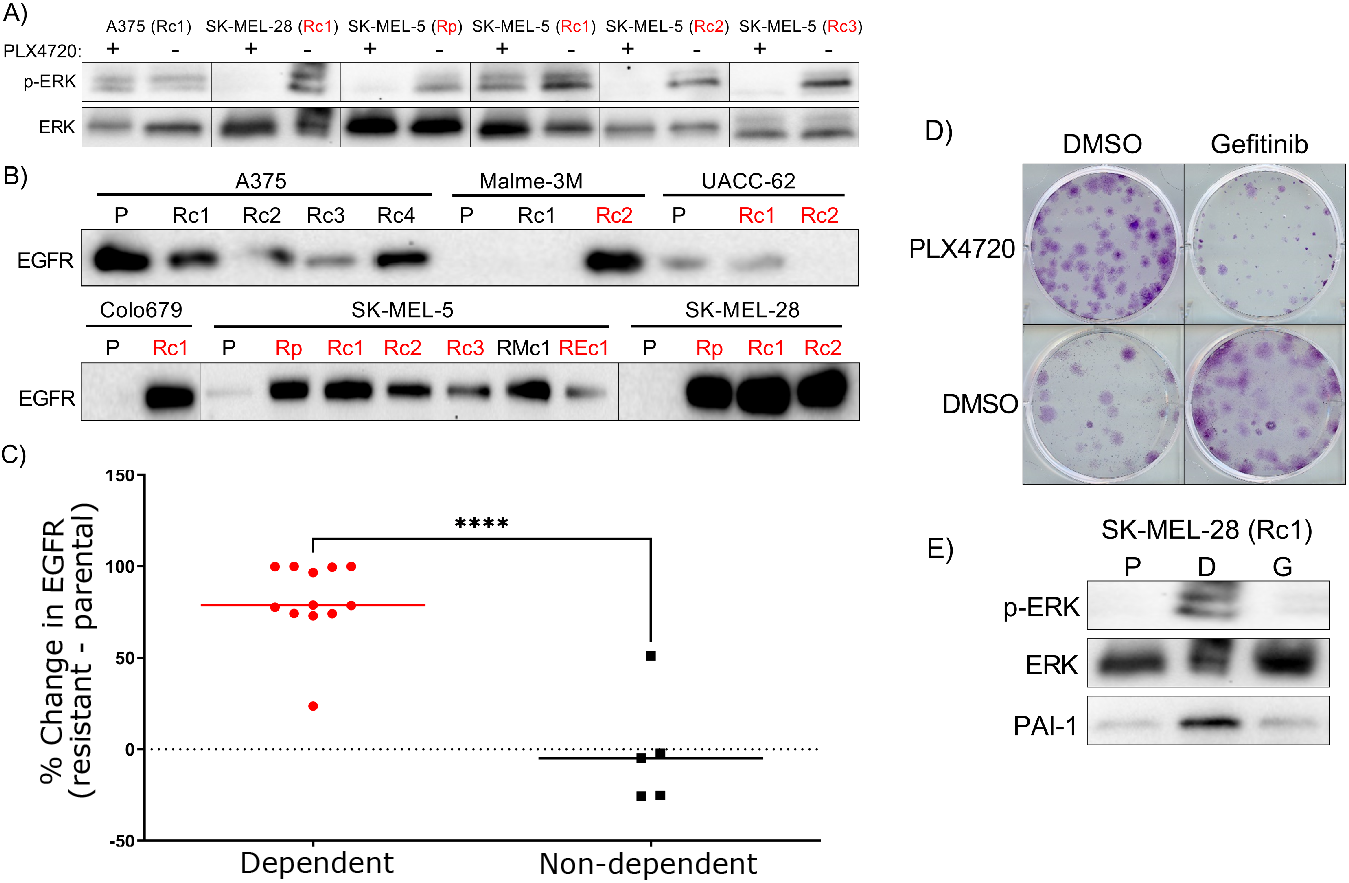
Dependent, but not non-dependent resistant cell lines upregulate EGFR upon resistance and have an increase of phospho-ERK on drug withdrawal. A) The indicated cells lines (R, PLX4720-resistant; c, clone, dependent cell lines highlighted in red) were treated with PLX4720 or DMSO for 4 hours and then immunoblotted for phospho-ERK and ERK. B) The indicated cell lines, dependent cell lines highlighted in red, were immunoblotted for EGFR. C) The immunoblot in B) was quantified and the percent change in EGFR expression relative to parental cell line was calculated. D) A375 PLX-resistant clone 4 was plated at low density and treated with DMSO, 3 μM PLX, 3 μM gefitinib and the combination the following day for 14 days. The resultant colonies were fixed and stained. E) SK-MEL-28 PLX-resistant clone 1 was treated with DMSO, 3 μM PLX or 3 μM gefitinib for 4 hours and immunoblotted for phospho-ERK and ERK. The same line was treated with the same conditions for three days and immunoblotted for PAI-1.

### Drug holidays can minimize growth when resistance is drug dependent

The poor growth of drug dependent resistant cells in drug-free environments suggests that population growth may be minimized by incorporating drug-free “holidays” into treatment schedules (70, 65). However, it is not clear how to design an optimal drug holiday regimen, which may depend on both the initial properties of the population (e.g. the ratio of sensitive to resistant cells) as well as the growth characteristics of different subpopulations. To investigate this question, we seeded a growth experiment with a mixed population of approximately 2 × 10^4^ cells comprised of 90 percent drug resistant cells–loosely similar to what one might find in a progressing tumor that has become dominated by resistant cells. In one population (SK-MEL-28), the resistant cells exhibited drug-dependence, while in the other population (A375), the resistant cells were not drug dependent. We then measured population size in each population after 9 days exposure to one of 3 different treatment protocols: 1) DMSO only each day, 2) inhibitor (PLX4720 at 1 *μ*M) only each day, or 3) an (arbitrary) periodic schedule consisting of drug treatments and drug holidays in a 2-1 temporal ratio. We found that in each population, the periodic dosing schedule outperformed either the drug-free or the drug-only treatments, but not both. Specifically, population size was minimized by drug-free treatment in the populations containing drug dependent (SK-MEL-28) resistant subpopulations and by drug-only treatment in the populations with non-dependent resistant subpopulations (Figure 4A-B). The relative ordering of the treatments is perhaps not surprising, given that the initial populations were dominated by drug resistant cells that favor either drug (drug dependent resistant) or nondrug environments (non-dependent resistance), but it raises the question of whether a more judiciously chosen treatment schedule may lead to reduced growth.

**FIG 4.**
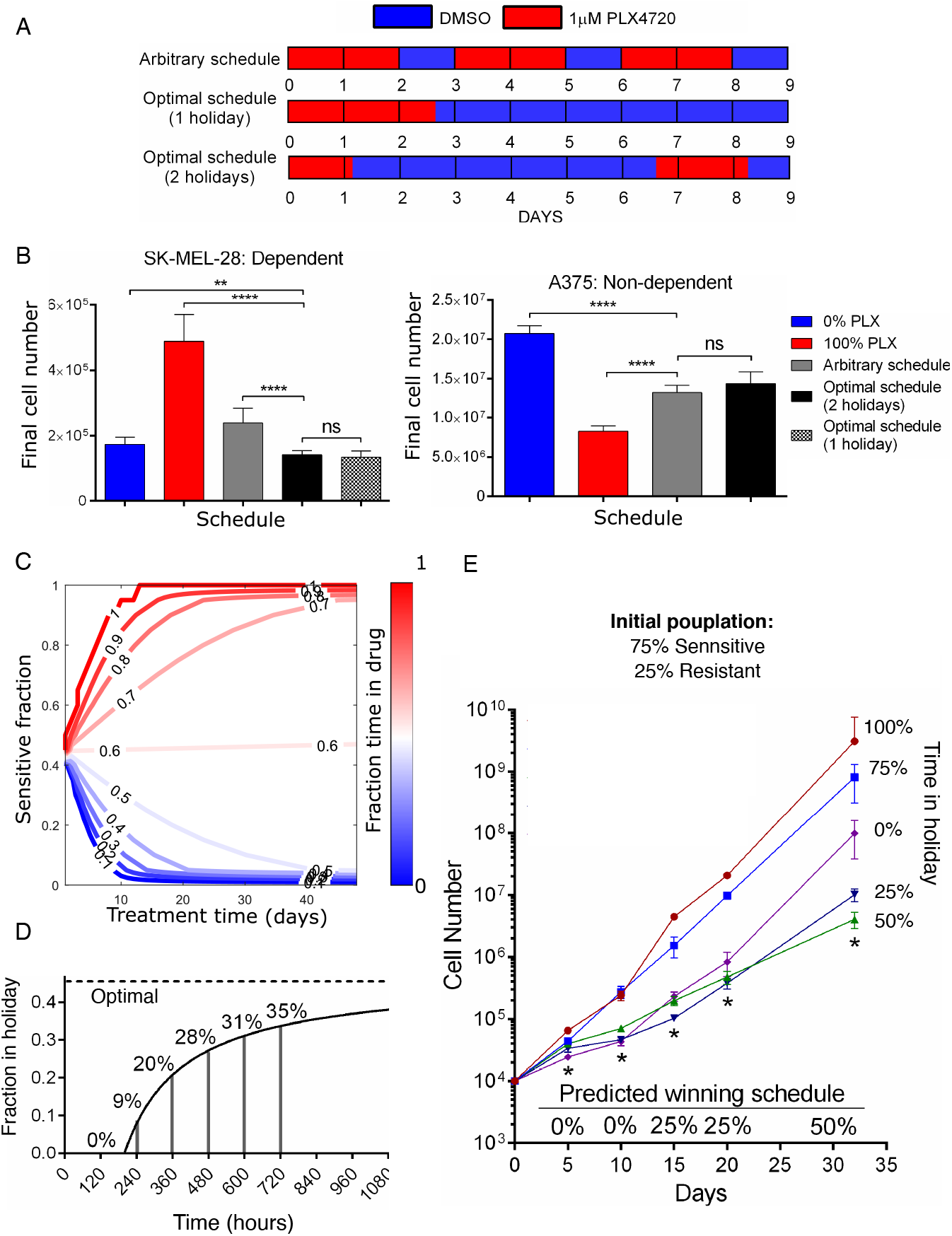
Scheduling drug exposure can optimize growth inhibition in cell lines that display dependency. A) A schematic of 5 different drug schedules: 1) drug-free (DMSO) every day; 2) drug (1 *μ*M PLX4720) every day; 3) arbitrary periodic schedule with epochs of drug and DMSO in a 2-1 temporal ratio; 4) model-inspired schedule with drug applied 29 percent of the time (up front); and 5) model-inspired schedule with drug applied 29 percent of the time but split into two epochs. B) Left panel: 2,000 SK-MEL-28 parental and 18,000 SK-MEL-28 PLX4720-resistant cells were plated in triplicate in 10 cm plates and treated according to the schematic in A, DMSO (0% PLX) or 1 μM PLX4720 (100% PLX) 24 hours later. On the ninth day, cells were trypsinized and counted. Right panel: 2,000 A375 parental and 18,000 A375 PLX4720-resistant cells were plated in triplicate per schedule and treated according to the schematic in A, DMSO (0% PLX) or 1 μM PLX4720 (100% PLX) 24 hours later. On the ninth day, cells were trypsinized and counted. C) Empirically constrained simulations show the optimal fraction of time exposed to drug as the ratio of sensitive to drug dependent cells and total treatment time is varied. D) Algorithm predicted optimal percent drug holiday (percent time in DMSO) as total treatment time increases. Optimal asymptote is represented with a dashed line. E) A mixture of 2,500 SK-MEL-28 parental and 7,500 SK-MEL-28 PLX4720-resistant cells were plated in triplicate and treated with the indicated schedule 24 hours later. Cells were trypsinized and counted every 5 days. The algorithm predicted optimal (‘winning’) schedule is indicated along the x-axis and time points that conform to predicted values are indicated with an asterisk.

To answer this question, we considered a simple mathematical model of exponentially growing subpopulations (sensitive and resistant cells) whose growth rates depend on the environment (with or without drug; see SI). On long timescales, we expect static treatments to eventually select for the subpopulation with the highest growth rate in that environment. However, in the presence of time-dependent treatments that switch between drug and no-drug epochs, it may be possible to maintain a heterogeneous population of both cell types–each one suboptimal in one environment, optimal in the other–that leads to reduced growth of the total population. Indeed, in the limiting case where cells do not interact, it is straightforward to show that a treatment strategy will maintain co-existing populations of both cell types when the drug treatments comprise a fraction (*f*_*d*_) of the total treatment time *T*,

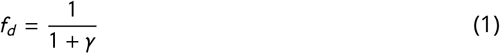

where 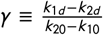 is a dimensionless parameter that depends only on the growth characteristics (*k*_*id*_) of the different subpopulations and *k*_*id*_ is the per capita growth rate of population *i* in drug concentration *d*. This simple model suggests that we can maintain population heterogeneity by incorporating drug-free epochs when *γ* > 0–that is, when sensitive cells grow faster than resistant cells without drug, while resistant cells grow faster than sensitive cells with drug. In the long-term limit (*T* → ∞), this treatment strategy yields a constant ratio of the different cell types, independent of the initial composition of the population. In addition, the growth of this heterogeneous population is optimal–that is, it is smaller than growth in either static with-drug or static drug-free environments–if

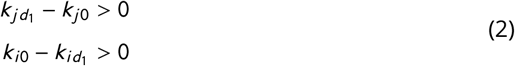

Equation 2 says that one cell type (*j*) must grow faster with drug than without, while the other cell type (*i*) must grow faster without drug than with–the precise conditions that define drug dependent resistance. In summary, in this simple model, one can maintain heterogeneity as long as sensitive cells are favored during drug-free epochs and resistant cells are favored during with-drug epochs. But that heterogeneity leads to minimal growth only when the resistant cells grow faster with drug than without (that is, when they are drug dependent).

Equation 1, along with experimentally measured growth rates of parental and resistant lines with and without drug, predicts an optimal schedule with *f*_*d*_ = 0.29 (holiday 71 percent of the time) for the SK-MEL-28 (drug dependent) population and *f*_*d*_ = 1 (drug-only) for the A375 (non-dependent) population. Based on these predictions, we experimentally tested two additional treatment schedules: 4) an optimal schedule with *f*_*d*_ ≈ 0.29 (2.6 days of drug exposure) where the holiday was applied at the end of the treatment, and 5) an optimal schedule with *f*_*d*_ ≈ 0.29 (2.6 days of drug exposure) where the holiday was applied over two different windows during the treatment. Consistent with predictions of the model, these schedules (4 and 5) outperformed other treatments in the populations with drug dependent resistance (SK-MEL-28), but were outperformed by the drug-only treatment (treatment 2) in the populations with non-dependent resistance (Figure 4A-B). We found no significant difference between schedules 4-5, consistent with the fact that schedules should depend only on *f*_*d*_ (not the specific timing of the holidays) when subpopulations are acting approximately independently.

### The optimal schedule is dependent on the treatment length

On sufficiently long time horizons, we expect that the optimal treatment will not depend on the initial composition of the tumor. However, many real-world applications are likely to involve finite-time treatments where these asymptotic results are not valid. Under these conditions, it is straightforward to show (see SI) that growth of the population will be minimized when

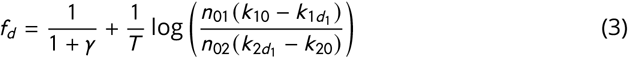

Equation 3 reduces to Equation 1 when *T* → ∞. However, on shorter time horizons, the two terms in Equation 3 may be comparable in size, and the optimal treatment will therefore depend on the total treatment duration *T* (Fig 4C). To test this prediction experimentally, we plated a mixed population of 10^5^ cells comprised of 25 percent resistant-dependent SK-MEL-28 cells. The cells were treated with one of five schedules with drug holidays ranging from 0 (static with-drug treatment) to 100 percent (static drug-free treatment). The treatments schedules were broken into 5 day blocks so that the cells could be counted every 5 days for an indeterminate length of time and remain on schedule. Based on results of the model (Fig 4D), we hypothesized that the treatment performing optimally would vary over time, with static drug treatments minimizing growth at early times while treatments with increasing drug holidays (asymptotically approaching the long-time optimal of approximately 45 percent drug holiday) minimize growth on longer timescales.

Indeed, experiments confirmed that growth was initially (e.g. days 5 and 10) minimized in the static drug (PLX4720) treatment (Fig 4E). However, by day 15, the treatment with 25 percent holiday was optimal, while the treatment with 50 percent drug holiday–which most closely corresponds to the long-time optimal–led to minimal growth by day 30. These results underscore the notion that the optimal strategy for drug holidays will initially vary over time–a phenomenon tied to the transient behavior of the population as it approaches a steady state composition–but for sufficiently long treatment periods approaches an optimal that is determined by the dimensionless parameter *γ* (and is therefore independent of the initial population composition).

### The addition of non-dependent resistant cells does not change the optimal treatment schedule

Because parental melanoma lines may give rise to both drug dependent and non-dependent resistant lineages, we next asked how the optimal treatment schedule would be impacted by the presence of both resistant types. To answer this question, we treated two different mixed populations containing drug sensitive, drug dependent resistant, and non-dependent resistant derivatives of UACC-62 with one of three treatments: 1) static drug-free treatment, 2) static treatment with PLX4720), or 3) a model-inspred (approximately) optimal treatment with *f*_*d*_ = 0.45 (55 percent of time in drug holiday). We chose UACC-62 cells for these studies simply because it is a line that gives rise to both resistant-dependent and resistant nondependent cells (Fig 1D). The two mixed populations were comprised of approximately constant ratios of drug sensitive to drug dependent resistant cells, but one population initially contained 0.2 percent non-dependent resistant cells and the other 10 percent non-dependent resistant cells. In both populations, the model inspired therapy– which contains drug holidays calculated without knowledge of the non-dependent populations–significantly outperformed the static therapies (Fig 5B). As one might intuitively expect, however, the population sizes are much larger in populations that started with more non-dependent resistant cells, consistent with predictions of the model (Fig 5A). This result highlights a more general principle: in many cases of interest, non-dependent (“traditional”) resistant cells grow at similar rates in the presence and absence of drug, at least over a range of drug concentrations. The presence of these non-dependent resistant cells is therefore not expected to alter the choice of optimal therapy (e.g. *f*_*r*_), but it can significantly limit the utility of such therapy in cases where these cells grow faster than the average growth rate of the drug-holiday optimized population of sensitive and drug dependent cells.

**FIG 5.**
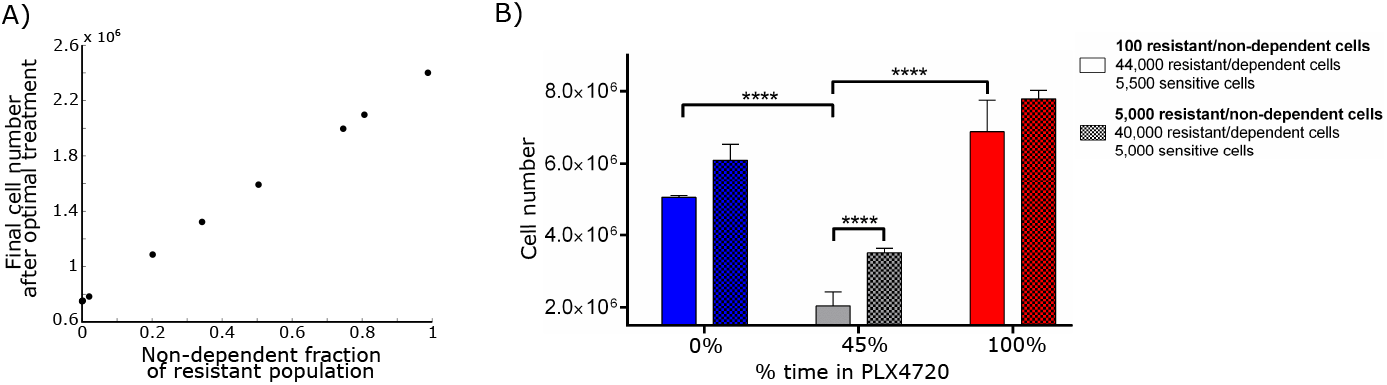
In a mix of sensitive, resistant non-dependent and resistant dependent cells, the fraction of dependent cells dictates the degree of benefit achieved by scheduling. A) Using empirically constrained simulations, we plot the total population count as the fraction of non-dependent resistant cells is varied from 0% to 100% of the resistant population. B) A mixture of 100 UACC-62 PLX4720-resistant clone 4 (resistant non-dependent), 44,000 UACC-62 PLX4720-resistant clone 3 and 5,500 UACC-62 parental cells or a mixture of 5,000 UACC-62 PLX4720-resistant clone 4 (resistant non-dependent), 40,000 UACC-62 PLX4720-resistant clone 3 and 5,000 UACC-62 parental cells were plated in triplicate and treated with the indicated schedules 24 hours later. Cells were trypsinized and counted on day 9.

### Adaptive therapy based on total population size can approximate optimal holiday schedules

While it is possible in the laboratory setting to measure the growth rates of each cell type with and without drug, in the clinical setting, such detailed characterization is not typically possible. Therefore, we attempted to develop an adaptive therapy method approximating the optimal holiday schedule in the absence of detailed growth rate information about the individual subpopulations (see Fig 6A, Methods). Briefly, we began by administering drug for one treatment window, followed by removal of the drug for one treatment window. The growth rates of each treatment window were recorded and the treatment that led to a lower growth rate was then continued. This treatment protocol was continued until the growth rate of the population surpasses that of the opposite treatment type, as recorded by the most recent treatment window of that type. At that time the treatment is switched. This process was continued until the treatment ended.

**FIG 6.**
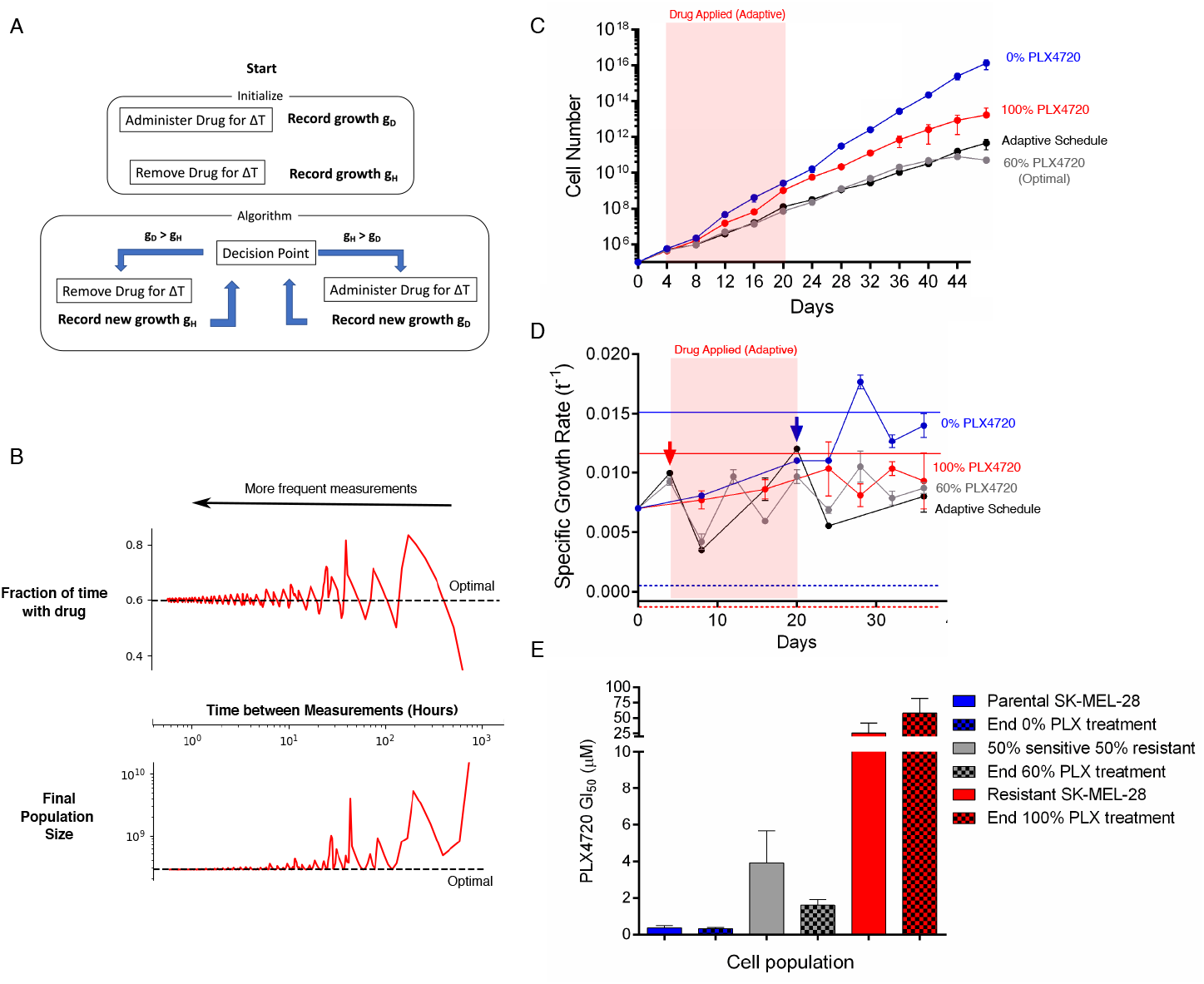
Optimal scheduling can be approximated with a mixed cell population of unknown growth rates. A) A cartoon schematic of the blind adaptive therapy algorithm. B) Top panel: Using empirically measured growth rates, we simulate the fraction of time spent exposed to drug following the blind adaptive therapy (solid red line) and compare it to the true optimal (dotted black line), as a function of how frequently population-level growth rates can be measured. Bottom panel: The same simulations as above, however instead of fraction of time spent in drug, the final cell count of the population (red solid line for blind therapy) is compared to what is projected from the optimal schedule (black dotted line). C) Results of the blind adaptive therapy experiment. Cell counts over time were recorded and the shaded red region highlights the period of time during which drug was applied. D) Growth rates for each schedule at the indicated time points. The growth rate of parental SK-MEL-28 cells in DMSO (blue solid line) and PLX4720 (blue dashed line) is indicated. Arrows indicate time points when the treatment condition was changed due to the measurement. The growth rate of PLX-resistant pooled SK-MEL-28 cells in PLX4720 (red solid line) and DMSO (red dashed line) is indicated. Red region highlights the period of time drug was applied. E) The PLX4720 GI50 values of the ending cell populations in E) as compared to the PLX4720 GI50 values of the indicated known mixtures of parental and resistant SK-MEL-28 cells.

We tested the performance of this approach computationally using our empirically measured growth rates from the previous experiments (Fig 6B). The success of this blind adaptive therapy treatment depends heavily on the frequency with which one could accurately measure total population growth. The simulations revealed that as the time between growth measurements shrinks, the blind adaptive therapy approaches the true optimal therapy schedule. In addition, the performance of the blind adaptive therapy schedule approaches that of the true optimal.

To test this approach experimentally, we established a mixed population comprised of sensitive and resistant dependent SK-MEL-28 cells and exposed replicate populations to one of four drug schedules: 1) static no-drug schedule, 2) static PLX4720 (constant drug) schedule, 3) the optimal schedule (*f*_*r*_ = 0.6, 40 percent drug holiday) based on measured growth rates of the different cell populations, and 4) the adaptive schedule described above. Treatments were applied in 4-day blocks for a total of 48 days. As predicted by the model, the optimal therapy (treatment 3) performed considerably better than the static with or without drug therapies. In addition, the adaptive therapy and the calculated optimal schedule performed almost identically over the course of 48 days (Fig 6C-D). The adaptive schedule achieves the goal of maintaining an average growth rate between the maximum and minimum growth rates for each cell type. In order to determine if we had achieved the goal of maintaining population heterogeneity, we took the final cell population from each schedule and measured the GI50 of each population to PLX4720 (Fig 6E). As a benchmark, we compared these GI50 values to those of freshly made mixes of 100% sensitive, 50% sensitive/50% resistant and 100% resistant populations. We found that the populations treated with 0% PLX4720 (100% PLX4720) exhibited GI50 values similar to those of fully sensitive (resistant) populations, consistent with the prediction that static therapies eventually lead to homogeneous populations (Fig 6E, blue and red bars). By contrast, populations treated with the optimal schedule exhibited GI50 values similar to, but slightly smaller than, those of an equally mixed population of 50% sensitive/50% resistant cells (Fig 6E, gray bars). Together, these results suggest it may be possible to reach near-optimal adaptive therapy treatment outcomes with incomplete population dynamics information.

## DISCUSSION

Our work complements a broad set of recent studies ranging from purely theoretical evolutionary therapy to clinical trials which have attempted to harness intermittent drug treatment schedules. Importantly, it demonstrates that while drug holidays can improve tumor growth control, they must be designed and implemented using strategies like the adaptive treatment regimen described here which account for the evolutionary dynamics of cells within the tumor harboring different drug response and dependency characteristics. The lack of these elements may be at least partially responsible for the failure of recent trials incorporating drug holidays. Further, this work also raises a number of new questions for future research. For example, having established that drug dependence varies dramatically with the choice of selecting inhibitor and cell line, it is clear that the genetic or cell state features of a tumor may make drug dependence more or less likely. Defining the mechanistic and evolutionary forces that lead to drug dependence, particularly across distinct cancers and targeted therapies, is an important step before treatment holidays can be optimally harnessed in the clinic. Finally, the notion that pharmacological manipulation of cell signaling or senescence may encourage the development of drug dependence is intriguing, as such therapeutic strategies have the potential to maximize the therapeutic value of adaptive drug holiday schedules.

It is also important to address several limitations in our approach. For example, we ignore the role of the host, specific resistance mutations, and ecological interactions within the tumor. The assumptions dramatically simplify the model, and though it provides a parsimonious description of these experiments, it also has features (e.g. independence of specific temporal ordering of treatments) that would likely break-down when intercellular interactions are included. In addition, these drug holiday experiments take place 1) in large populations where demographic stochasticity plays a minimal role and 2) on short timescales were additional evolutionary adaptation is likely negligible. Incorporating stochastic events, such as extinction and mutation, will likely extend the scope of the models and may uncover additional dynamics.

Yet even in this well controlled laboratory setting, calculating an accurate optimal schedule is challenging, and it is notable that such a simple mathematical model captures the main qualitative features of the dynamics. These simplifications make it impossible to use our model is not directly applicable in the clinic. Instead, our work is meant as a launching point in understanding how one might optimally treat a population containing drug dependent cells. Our results are significant because they show that optimization is possible, given a robust experimental setup and proper treatment of the underlying variables. Our hope is that future work might address these challenges and extend these basic principles for optimal scheduling to realistic clinical scenarios. In the long run, holiday-based schedules are a promising therapeutic avenue that may eventually lead to improved treatment outcomes for a subset of patients with drug dependent metastatic disease.

## MATERIALS AND METHODS

### Cell lines

A375, Colo679, UACC-62 and Malme-3M cells were grown in RPMI 1640 (Life Technologies Corporation, Carlsbad, CA) supplemented with 10% fetal bovine serum (Sigma-Aldrich Corporation, St. Louis, MO) and 1% penicillin/streptomycin (Life Technologies Corporation). SK-MEL-28 and SK-MEL-5 cells were cultured in Dulbecco’s modified Eagle’s medium (DMEM) (Life Technologies Corporation) with 10% fetal bovine serum and 1% penicillin/streptomycin. Malme-3M and SK-MEL-28 cell lines were obtained from L. Garraway (Harvard University, Dana-Farber Cancer Institute). All other cell lines were purchased from the American Type Culture Collection. All lines were submitted to STR profiling by the Duke University DNA Analysis Facility to confirm their authenticity.

### Chemicals

All inhibitors were purchased from Selleck Chemicals (Houston, TX) and prepared at 100 mM stock solutions in DMSO.

### GI_50_ assay

Wherever GI_50_ values of specific inhibitors are indicated, they were determined as follows. Cells were trypsinized and seeded at 5,000 cells/well in 96-well plates in normal tissue culture conditions. After a 24-hour incubation, diluent (typically DMSO) or concentrated 10-fold dilutions of the indicated inhibitors (at 1:1000) were added to the cells to yield a highest concentration of 40 μM (VX-11E) or 200 μM (all other inhibitors). After a three-day incubation with the treatment, cell viability was assessed with the CellTiter-Glo luminescent viability assay (Promega Corporation, Durham, NC) according to manufacturer’s instructions. Growth inhibition was calculated as a percentage of diluent-treated cells and GI50 values were determined to correspond to the inhibitor concentration that resulted in half-maximal growth inhibition.

### Immunoblotting

In order to measure protein levels in whole cell lysates, aliquots of cell extracts prepared in RIPA lysis buffer (FORMULATION (Sigma-Aldrich)) were submitted to SDS-PAGE. After electrophoretic transfer to PDMF, filters were blocked in 5% BSA and probed overnight at 4°C with the following primary antibodies and dilutions: p16 (1:200, #554079 BD Biosciences, San Jose, CA), p53 (1:200, #VMA00019 Thermo Fisher), phospho-Akt (Serine 473; 1:1000, #4058 Cell Signaling Technology, Danvers, MA), AKT (1:1000, #4691 Cell Signaling Technology), phospho-ERK (1:1000, #9101 Cell Signaling Technology), ERK (1:1000, #4695 Cell Signaling Technology), EGFR (1:1000, #2232 Cell Signaling Technology), PAI-1 (1:500, #11907 Cell Signaling Technology), PARP (1:1000, #9542 Cell Signaling Technology), vinculin (1:500; #4650 Cell Signaling Technology), α-tubulin (1:1000, #2125 Cell Signaling Technology) and β-actin (1:1000, #4970 Cell Signaling Technology). After washing in TBS-T, filters were incubated for 1 hour at room temperature with alkaline phosphatase-coupled goat antirabbit antibodies and developed with Western Lightning Plus (Perkin Elmer, Inc., Waltham, MA) according to manufacturer directions.

### Lentivirus preparation and DNA constructs

All expression clones were prepared in lentiviral form as previously described (75). In brief, vectors were packaged in 293T cells with an overnight incubation with Fugene (Promega Corporation), psPAX2 and pVSV-G. The virus-containing media was collected after 48 and 72 hours and filtered with a 0.45 μm filter and stored at −80°C until use with 16 μg/mL polybrene (Sigma-Aldrich). The sgRNA constructs used are listed in Table S2.

### In vitro adaption of inhibitor resistant cells

PLX4720-, AZD6244-or VX-11E-resistant cell lines were produced by one of two methods as previously described (75). Briefly, parental cells were either exposed to escalating doses of inhibitor until logarithmic growth resumed or exposed to a high dose (3 μM) of inhibitor and the resultant resistant clones were expanded and cultured. Parental cell lines were cultured concurrently with DMSO. Resistant cell lines were maintained in routine culture with the addition of 3 μM inhibitor. All resistant and DMSO parental control lines were submitted to STR profiling by the Duke University DNA Analysis Facility upon the acquisition of resistance in order to confirm their authenticity.

### Clonogenic growth assay

To measure the ability of cell lines to form colonies from a single cell, clonogenic growth assays were performed as previously described (76). Briefly, 100 cells were seeded per well in 6 or 12-well tissue culture plates in normal growth media. After 24 hours, the indicated treatments were added and the assay was incubated for 14 days with the addition of fresh media and inhibitors after 7 days. Cells were fixed and stained with 0.5% (w/v) crystal violet in 6.0% (v/v) glutaraldehyde (Thermo Fisher Scientific, Waltham, MA).

### Annexin V apoptosis assay

The induction of apoptosis was quantified as described previously (Martz et al., 2014). Briefly, cells were plated in triplicate at 200,000 cells per well in six-well plates. The following day, the growth media was removed and replaced with fresh media containing the indicated dose of drug or diluent (typically DMSO). After a 72-hour incubation in drug, cells were washed in PBS twice and resuspended in a buffer composed of 10 mM HEPES, 140 mM NaCl and 2.5 mM CaCl2 (BD Biosciences, San Jose, CA). Apoptosis was quantified using allophycocyanin-conjugated Annexin V and viability was assessed with 7-Amino-actinomycin D (BD Bioscience). Gating was defined using untreated/unstained cells and treatments were evaluated at 20,000 counts using BD FACSVantage SE.

### Cell cycle analysis

In order to analysis the progression of cells through the cell cycle, cells were plated in triplicate at 200,000 cells per well in six-well plates. The following day, the growth media was removed and replaced with fresh media containing the indicated dose of drug or diluent (typically DMSO). After a 72-hour incubation in drug, cells were trypsinized and counted. 1×106 cells were washed twice with PBS and then fixed in 70% ethanol. The cells were washed twice in PBS and stained with a solution of 50 μg/μL RNase A, 20 μg/μL propidium iodine, 0.05% TritonX-100 in PBS (Sigma-Aldrich). DNA content of the cell population was determined and quantified by flow cytrometry. Gating was defined using untreated/unstained cells and treatments were evaluated at 20,000 counts using BD FACSVantage SE.

### Senescence-associated *β*-galactasidase assay

In order to determine the effect of treatment on the induction of senescence, cells were plated in triplicate in 6-well plates at a density that would achieve 80-90% confluency at 10 days of growth, typically between 1,000 and 50,000 cells. After 24 hours of culture, the growth media was removed and replaces with fresh media containing the indicated drug or diluent. After 10 days of treatment, media was removed and the cells were stained with the kit, Senescence β-Galactosidase Staining Kit (Cell Signaling Technology), according to manufacturer’s instructions. Bright field images were taken at 100X magnification at five random locations in each well. Representative images are shown.

### Growth rate calculation

To calculate growth rates, cell populations were used at low passage number with measured drug resistance. Cells were plated in triplicate in 10 cm plates at 3,000 cells per plate in normal growth media. The following day, cells were treated with drug or diluent. Six days later, the cells were lifted with 0.25% trypsin (Life Technologies) and counted using a Z2 Coulter Particle Count and Size Analyzer (Beckman Coulter, Pasadena, CA). Growth rates (μ) were calculated from the number of cells plated (N0) and the number counted (N) according to the formula:

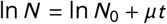

where t is the elapsed time, or 168 hours.

### Drug scheduling assay

To test the effects on cell number by a drug schedule, the following was performed. Immediately after growth rates were measured, the indicated number of cells were plated in growth media with no drug in 15 cm plates. The following day, the indicated drug schedule was added to the cells. Cells were trypsinized and counted at the end of each completed schedule and replated as indicated. All drug schedules were establish based on the interval between counts, for example, if cells were treated with 75% drug and counted every four days, they would be treated with drug for three days and diluent for one day, counted, and replated for the duration of the experiment. If a portion of cells was discarded before replating, growth rates (μ) were calculated over the period from the number of cells plated at the beginning of the period (N0) and the number counted at the end of the period (N) according to the formula:

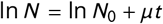

where t is the period length in hours. These growth rates were then used to project total cell number as if no cells had been discarded. After the initial plating, which was always into diluent only, cells were replated directly into the treatment determined by their schedule.

### Statistics

All results are shown as means±SD. In order to compare groups, unless noted otherwise, P values were determined using unpaired, two-tailed Student’s t tests.

### “Blind” adaptive therapy algorithm

To begin, administer drug for some time Δ*T* and record the growth rate during that period. This period is followed by the removal of drug for the same Δ*T* and the drug-free (or ‘holiday’) growth rate is recorded. Administer the condition with the smaller growth rate and record the growth rate for each Δ*T* that passes. Decision point: if the growth rate for the most recent Δ*T* surpasses that of the opposite condition, switch the treatment to the opposite condition for the next time window. Repeat this process for the rest of the treatment time continually recording new values for the population growth in drug and during treatment holiday and switching between treatments as they become optimal.

## ACKNOWLEDGMENTS

This work was supported by Duke University School of Medicine start-up funds (to KCW), the Duke Cancer Institute (to KCW), and NIH awards R01 CA207083 (to KCW), R01 CA263593 (to KCW), and NIH R35 GM124875 (to KBW). The format for this preprint is adapted from the ASM template (mSystems) available on Overleaf.com.

