## Supplemental Text for "Drug dependence in cancer is exploitable by optimally constructed treatment holidays"

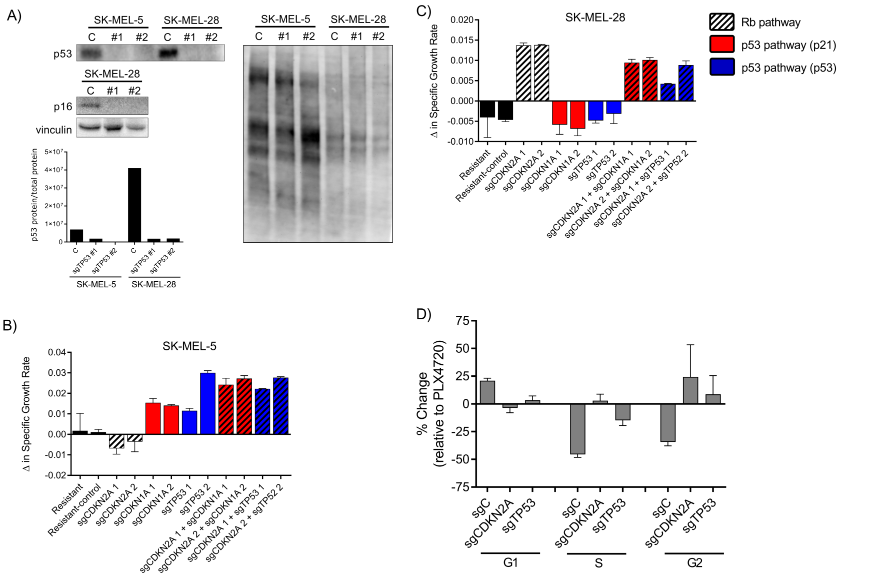


**FIG S1** **Drug dependence is associated with drug withdrawal-induced senescence.** A) SK-MEL-28 and SK-MEL-5 PLX-resistant cells expressing sgControl or 1 of 2 sgTP53 constructs (top left panel) and SK-MEL-28 PLX-resistant cells expressing sgControl or 1 of 2 sgCDKN2A constructs (bottom left panel) were treated with PLX4720 withdrawal for three days and immunoblotted for p53 or p16. Total protein (right panel) and quantification (bottom panel) are shown. B) Growth rates of the indicated cell lines were measured over 7 days of PLX4720 or DMSO treatment. The change in growth rate in DMSO relative to PLX4720 treatment is shown. C) As in B), in the SK-MEL-28 PLX-resistant cells. D) SK-MEL-28 PLX4720-resistant cells expressing the indicated constructs were treated with DMSO or PLX4720 for 3 days and cell cycle analysis was performed as described in the Material and Methods.

### A SIMPLE MODEL OF $N$ SUBPOPULATIONS

Consider a simple model with $N$ different cell populations. The $i^{\text{th}}$ cell population grows exponentially with per capita growth rate $k_{id}$ in the presence of drug at concentration $d$. The entire metapopulation evolves according to

$$n = G_d n, \tag{1}$$

where $n$ is a vector with components $n_i(t)$ (the number of cells of type $i$ at time $t$) and $G_d$ is an $N$-by-$N$ square matrix whose diagonal entries are $k_{id}$ for $i = 1, 2, \ldots N$ and whose off-diagonal entries correspond to interconversions between different cell types. Without loss of generality, we can rescale time and measure rate constants in units of $k_{10}$ (the growth rate of population $i = 1$ in the absence of drug). The solution to Equation 1 is given by $n(t) = A_d(t)n(0)$, where the $A_d(t) \equiv e^{G_d t}$ is a time dependent matrix and $e$ denotes the standard matrix exponential. Similarly, if the cell population is exposed to a treatment of duration $t_0$ with drug at concentration $d_0 = 0$ followed by a treatment of duration $t_1$ with drug at concentration $d_1 > 0$ (with $T \equiv t_1 + t_2$), the solution is given by

$$n(T) = M(t_1, t_2)n(0), \tag{2}$$

where $M(t_1, t_2) \equiv A_d(t_1)A_0(t_0)$. If we imagine repeating the periodic dosing protocol a total of $q$ times, the solution is simply

$$n(qT) = M^q n(0), \tag{3}$$

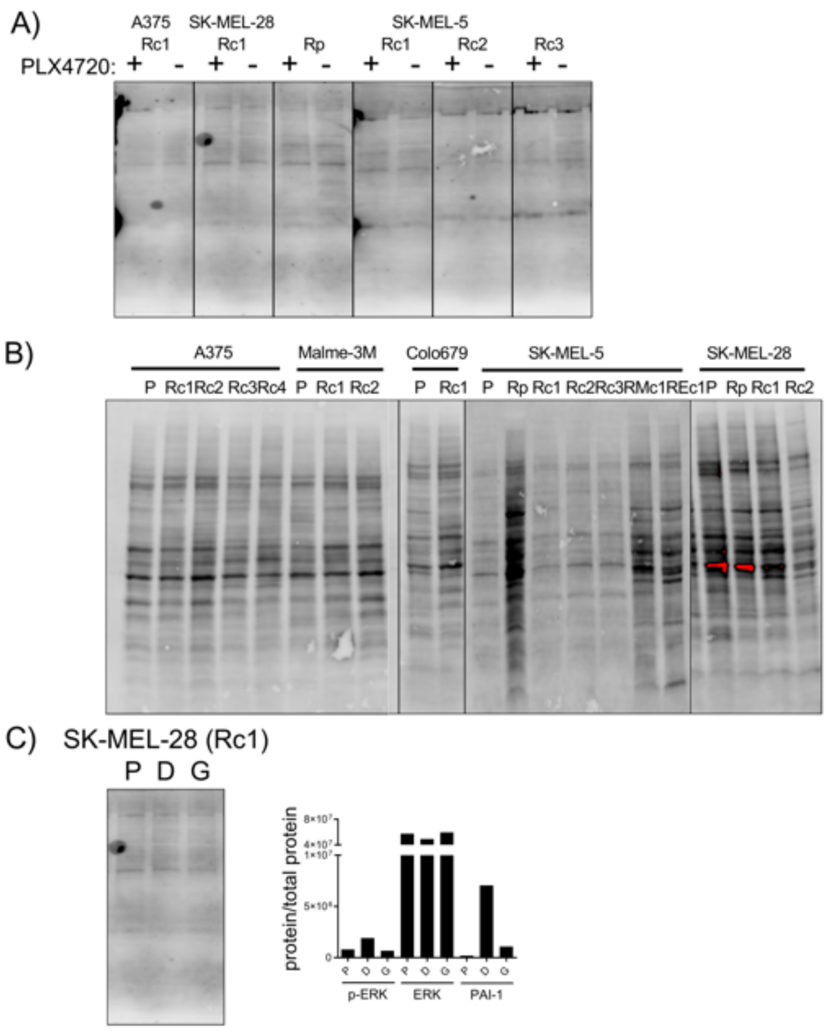


**FIG S2** Protein loading quantification corresponding to immunoblots in Fig. 3. A) Total protein blots for the immunoblots shown in Fig. 3A. B) Total protein blots for the immunoblots shown in Fig. 3B. C) Total protein blots (left panel) and quantification (right panel) for the immunoblots shown in Fig. 3E.

where we've dropped the explicit dependence of $M$ on $t_1$ and $t_2$ for economy of notation.

**Long-time solution.** We are primarily interested in the long-time behavior of Equation 3; that is, we are interested in solutions in the limit $q >> 1$ such that the periodic dosing protocol has been applied many times. In this regime, the growth will be dominated by the largest eigenvalue of $M$, which we denote by $\lambda_{\max}$, and the population will grow exponentially at a per-capita growth rate of $g$ given by (77)

$$g = \frac{\ln \lambda_{\max}}{T}. \tag{4}$$

The long-term population will be comprised of $j$ sub-populations when the dominant eigenvector of $M$ has $j$ nonzero entries.

**Wild-type and Mutants.** Consider now a case where population $i = 1$ corresponds to a "wild-type" progenitor cell. These cells can mutate to each of the remaining $N - 1$ cell types ("mutants") with some rate $\epsilon$. We consider these mutations to be irreversible and neglect reversion to wild-type cells or interconversion between different mutant cells. Hence, the matrix $G_d$ is given by

$$(G_d)_{i,j} = \delta_{ij}(k_{id} - \delta_{11}(N-2)\epsilon) + \delta_{i1}\epsilon, \tag{5}$$

where $\delta_{ij}$ is the usual Kronecker delta. In the limit $\epsilon \to 0$, the matrix $G_d$ (and in turn, the matrix $M$) is diagonal with off-diagonal terms of order $\epsilon$. If we assume $\epsilon \to 0$, $G_d$ is a diagonal matrix whose (i,i)$^{\text{th}}$ entry is $k_{id}$. In turn, the matrix $M$ is also diagonal with eigenvalues given by

$$\lambda_i = e^{k_{i0}t_0 + k_{id_1}t_1} = e^{T(k_{i0}(1-f_d)+k_{id_1}f_d)}, \tag{6}$$

where $f_d \equiv t_1/T$ is the fraction of time spent in drug, and eigenvectors are given by the standard basis vectors for $\mathbb{R}^N$. The different cell populations are uncoupled, and the dominant population is the one with the largest value of $\lambda_i$. In this limit, the temporal ordering of drug and no-drug regimens does not matter; the outcome is determined entirely by $f_d$, the fraction of time spent in drug.

**Coexistence of subpopulations.** If we choose $t_0 = 0$ (drug always present) or $t_1 = 0$ (drug never present), the population will eventually be dominated by the cell type with the fastest per capita growth rate in the presence or absence of drug, respectively. On the other hand, there may be cases where allowing multiple cell types to co-exist decreases the total population growth. To maintain coexistence of two cell types, $i$ and $j$, one requires that $\lambda_i = \lambda_j$, which yields

$$(k_{i0} - k_{j0})(1 - f_d) + (k_{id_1} - k_{jd_1})f_d = 0. \tag{7}$$

Solving for $f_d$, we have

$$f_d = \frac{1}{1+\gamma}, \tag{8}$$

with $\gamma \equiv \frac{k_{id_1} - k_{jd_1}}{k_{j0} - k_{i0}}$. Note that $0 < f_d < 1$ if and only if $\gamma > 0$ and finite. In other words, one cell type must be favored in the presence of drug, while the other cell type must be favored in the absence of drug. Choosing $t_1 = f_d T$ will conserve the ratio of cell types $i$ and $j$, and the long-term per capita growth of the population is

$$g = \frac{k_{i0}k_{jd_1} - k_{j0}k_{id_1}}{(k_{i0} - k_{id_1}) - (k_{j0} - k_{jd_1})}. \tag{9}$$

It is not possible, in general, to choose $f_d$ such that more than two cell types coexist asymptotically. Doing so would require $\lambda_i = \lambda_j = \lambda_k$, or equivalently,

$$Kt = 0, \tag{10}$$

where $t$ is a column vector with entries $t = (t_0, t_1)$ and $K$ is a matrix of growth rate differences given by

$$K \equiv \begin{pmatrix} k_{i0} - k_{j0} & k_{id_1} - k_{jd_1} \\ k_{i0} - k_{k0} & k_{id_1} - k_{kd_1} \end{pmatrix}. \tag{11}$$

Equation 10 has a nontrivial solution only when $\det K = 0$–that is, a non-trivial solution exists only when the growth rates satisfy a particular condition (specifically, when the columns of $K$ are linearly dependent). Aside from this special case, it is not possible to maintain co-existence between more than two cell types in the long time limit.

Similar results also apply if we consider cell type $i = 1$ (wild-type) and take $N\epsilon << 1$ but non-zero. In that case, the eigenvalue corresponding to wild-type cells is given by Equation 6 with $k_{i0} \to k_{i0} - (N-1)\epsilon$ and $k_{id_1} \to k_{id_1} - (N-1)\epsilon$. The other eigenvalues remain unchanged. Similarly, the eigenvector corresponding to wild-type cells is given by $v = (1 - O(\epsilon), b_1, b_2, ..., b_N)$, where $b_i \sim O(\epsilon)$. In words, this means that when $f_d$ is chosen such that $\lambda_1$ dominates, the long-term population maintains a small fraction ($O(\epsilon)$) of all mutants. On the other hand, if $f_d$ is chosen such that $\lambda_1$ and $\lambda_j$ dominate, the population will consist of co-existing populations of wild-type cells and type $j$ cells as well as a diminishing fraction ($O(\epsilon)$) of all additional mutants. Of course, as $\epsilon$ approaches the size of the other growth rates in the problem, mutation (and cell interconversion more generally) can have a significant effect on the overall dynamics (see, for example, (78)).

**Optimality of coexisting subpopulations.** It is possible to maintain heterogeneity in a population as long as $\gamma > 0$. When is it optimal to adopt such a periodic dosing strategy? That is, when does this heterogeneous population exhibit a lower average growth rate than each of the corresponding homogeneous populations? To answer this question, we first assume $\gamma > 0$ such that the long-term population is heterogeneous. Without loss of generality, we consider a population consisting of two subpopulations $i$ and $j$ and take $k_{i0} > k_{j0}$ and $k_{jd_1} > k_{id1}$ to ensure $\gamma > 0$. That is, we assume that population $i$ grows faster than $j$ in the absence of drug and $j$ grows faster than $i$ in the presence of drug. In order for this heterogeneous population to represent the optimal (slowest-growing) population, we require that $g$ (given by Equation 9) is smaller than the per capita growth rate of the fastest growing population in each condition alone. Specifically, we have

$$g < k_{i0} \text{ and } g < k_{jd_1} \tag{12}$$

By combining Equation 9 with Equation 12, we have

$$\begin{aligned} \frac{(k_{jd_1} - k_{id_1})(k_{jd_1} - k_{j0})}{(k_{i0} - k_{id_1}) - (k_{j0} - k_{jd_1})} &> 0, \\ \frac{(k_{i0} - k_{id_1})(k_{i0} - k_{j0})}{(k_{i0} - k_{id_1}) - (k_{j0} - k_{jd_1})} &> 0. \end{aligned} \tag{13}$$

Excluding the singular case where the denominator is zero, we are therefore led to two optimality conditions:

$$\begin{aligned} k_{jd_1} - k_{j0} &> 0 \\ k_{i0} - k_{id_1} &> 0 \end{aligned} \tag{14}$$

In words, cell type $j$ must grow faster with drug than without, while the opposite must be true for cell type $i$. Hence, for population heterogeneity to be optimal in the long-time limit, we must have one subpopulation of drug-sensitive cells and one population of drug-addicted cells.

**Exact solution for N=2.** In the simple case where $N = 2$ and $\epsilon = 0$, it is possible to write down an analytic expression for the fractional time in drug ($f_d$) required to minimize population size. The expression is valid for all times, not merely in the long time limit. To do so, consider that the total population size at the end of period $T$ is governed by Equation 2. Because $M$ is diagonal, one can trivially write the solution for $P = n_1 + n_2$ as

$$P = n_{01} \exp\left(T(k_{10}(1 - f_d) + k_{1d_1} f_d)\right) + n_{02} \exp\left(T(k_{20}(1 - f_d) + k_{2d_1} f_d)\right), \tag{15}$$

where $n_{0i}$ is the initial size of population $i$. It is straightforward to show that $\partial^2 P/\partial f_d^2 < 0$, so this is a convex function with a unique minimum on $f_d = [0, 1]$. One can solve for the minimum according to $\partial P/\partial f_d = 0$, which gives

$$f_d = \frac{1}{1+\gamma} + \frac{1}{T} \log\left(\frac{n_{01}(k_{10} - k_{1d_1})}{n_{02}(k_{2d_1} - k_{20})}\right) \tag{16}$$

with $\gamma \equiv \frac{k_{1d_1}-k_{2d_1}}{k_{20}-k_{10}}$. In the long-time limit ($T \to \infty$), Equation 16 reduces to Equation 8. However, when the addiction criteria (Equation 14) is not met, the second term in Equation 16 is imaginary. As a result, the long-time limit (Equation 8) corresponds to an optimal solution only when $T = \infty$; for any finite time, the long-time limit will, in general, not be optimal and cycling is not beneficial, as $f_d$ is complex with a nonzero imaginary part. On the other hand, when the addiction criteria is satisfied, the solution will converge continuously to the long-time limit.

**Summary of Cycling, Heterogeneity, and Optimality.** To maintain population heterogeneity in the long-time limit, one must cycle between drug and no-drug conditions with $f_d$ (the time spend in drug) given by Equation 8. In practice, one cell type must be favored in the presence of drug and the other cell type must be favored in the absence of drug. If $f_d$ does not satisfy Equation 8, the population will eventually be dominated by one population or the other. Specifically, it will be dominated by the population corresponding to the largest eigenvalue $\lambda_i$, even if a cycling protocol is used. It is not, in general, possible to asymptotically maintain finite fractions of more than 2 cell types by switching between 2 environments. The exception occurs when $K$ (Equation 11) has linearly dependent columns. In the limit where cells do not interact or interconvert, the total fraction of time spent in drug ($f_d$) determines the outcome, independent of the temporal ordering of the drug and no-drug regimens.

Maintaining heterogeneity through cycling is optimal only when one cell type grows faster with drug than without ("addiction"), while the other cell type grows faster without drug than with drug ("drug sensitive"). It is worth noting that it is possible to maintain population heterogeneity but *not* achieve optimality. This situation occurs when the above condition for optimality is not met, but $0 < f_d < 1$ is chosen according to Equation 8. In this situation, the average growth of the heterogeneous population is larger than the growth of the dominating population in at least one of the two conditions (either with or without drug). Of course, it is also possible under some conditions to achieve optimality without maintaining heterogeneity. This situation occurs, for example, when $\gamma < 0$, meaning that one cell type is favored in both conditions. In that case, one simply chooses the condition in which that cell type grows most slowly. It can also occur when $\gamma > 0$ but addiction is not present (e.g. mixture of a drug sensitive population and a drug-resistant, but not addicted, population with a small fitness cost). In that case, the mutant can be favored over the sensitive cells in the presence of drug, while the wild-type can be favored over mutant in the absence of drug. However, both cell types grow more slowly in drug. In that case, it is possible to maintain both cell types, but the optimal solution is to apply drug always, eventually leading to only resistant mutants.
